# DIOPT: the DRSC Integrative Ortholog Prediction Tool, 2026 update

**DOI:** 10.64898/2026.04.15.718708

**Authors:** Yanhui Hu, Aram Comjean, Chenxi Gao, Austin Veal, Shinya Yamamoto, Stephanie E. Mohr, Norbert Perrimon

## Abstract

Mapping orthologous proteins is a critical step for cross-species literature mining, data integration, experimental design, and more, making the ability to quickly predict orthologs across species a key tool for functional genomic studies. The DRSC Integrative Ortholog Prediction Tool (DIOPT) was initially developed in 2011 to provide a centralized portal for identifying inferred orthologs among major model organisms. By integrating results from multiple ortholog prediction algorithms, DIOPT allows users to compare predictions across methods and prioritize high-confidence ortholog relationships. Over the years, we regularly updated the underlying genome annotations and refreshed predictions from each integrated algorithm. In addition, both the number of supported species and the number of ortholog prediction algorithms incorporated into the platform have grown. The web portal has also been enhanced with new features designed to improve usability, facilitate data exploration, and support a broader range of research applications. We also developed a sister version of DIOPT tailored specifically for arthropod species; this enables researchers working with a diverse set of insects and related organisms to perform ortholog mapping and comparative analyses more effectively. Together, these developments ensure that DIOPT remains a robust and broadly useful resource for functional genomics research.

## INTRODUCTION

Orthologs are evolutionarily related genes in different species derived from the same gene in the last common ancestor, and tend to encode proteins with similar functions at a biochemical, cellular, and organismal level. By contrast, paralogs are genes found within the same species that originated by duplication of a single ancestral gene, and might have evolved new or divergent functions over time (Fitch 1970). Many research groups have developed sophisticated approaches for identifying orthologs, including methods based on the evolutionary history of genes and genomes (Emms and Kelly 2019; Fuentes *et al*. 2022; Altenhoff *et al*. 2024), the primary amino acid sequences of proteins encoded by genes (Walsh *et al*. 1987; Persson and Sonnhammer 2023; Cosentino *et al*. 2024), and/or protein domain organization (Persson *et al*. 2019). No single method for predicting ortholog relationships is the ‘right’ or ‘complete’ method, as no method has both perfect sensitivity, i.e., detecting all possible orthologs, and perfect specificity, i.e., excluding any pairs that are not true orthologs. Therefore, in 2011, we launched an integrative resource that combines prediction results from multiple resources to increase sensitivity along with a series filters that help users retrieve orthologs at various levels of specificity (Hu *et al*. 2011). The number of users of the DIOPT online tool has steadily grown since then, and DIOPT-based ortholog predictions have been integrated at other sites (e.g., the Alliance of Genome Resources and MARRVEL)(Wang *et al*. 2017; ALLIANCE OF GENOME RESOURCES 2022; ALLIANCE OF GENOME RESOURCES 2024).

Similar resources include HCOP (Wright *et al*. 2005; Yates *et al*. 2021) and OrthoList (Shaye and Greenwald 2011; Kim *et al*. 2018). HCOP initially integrated results from six ortholog mapping algorithms/resources; currently it includes results from 12. The choice of integrated resources at HCOP is similar to DIOPT. However, unlike DIOPT, which provides pair-wise mapping for all species covered, HCOP is human centric by design, providing only mapping between human genes and genes from other organisms. OrthoList was launched in 2011, integrates four algorithms/resources, and provides only mapping from *C. elegans* to human genes. In addition, compared with similar resources, the DIOPT online resource provides more features, including the option to view a protein alignment, search for paralogs within a species, and do one-to-all-species ortholog searches. The current version of DIOPT (v10) supports ortholog mapping among 13 species: *Homo sapiens, Mus musculus, Rattus norvegicus, Xenopus tropicalis, Danio rerio, Caenorhabditis elegans, Drosophila melanogaster, Anopheles gambiae, Ixodes scapularis, Arabidopsis thaliana, Schizosaccharomyces pombe, Saccharomyces cerevisiae*, and *Escherichia coli*, and integrates ortholog mapping from 19 different ortholog algorithms or resources.

To meet requests from the community not met by the online portal, we developed a standalone pipeline for customized orthologous mapping with user-specified choice of species and methods. This pipeline can be used to include species not currently covered by the DIOPT database as well as to make customized assemblies based on the subset of tools specified by the user. In addition, to further expand DIOPT for non-model organisms, we developed a sister database, DIOPT Arthropod Plus, focused on arthropods relevant to food security (i.e., crop pests) and human disease (i.e., vectors like mosquitos and ticks). *Drosophila melanogaster* serves as a “reference insect” for DIOPT Arthropod Plus because of the high quality and depth of gene annotations available for this species. The new DIOPT resource enables researchers to rapidly interpret and test mechanisms in a wide range of less tractable insect species. For example, the discoveries in flies can help prioritize candidate genes and pathways in other arthropods, including those involved in pesticide resistance, host preference, diapause, or virus susceptibility, therefore making downstream experiments in non-model insects more targeted and efficient.

## MATERIAL AND METHODS

### 1. Data sources and algorithms for DIOPT and DIOPT Arthropods Plus

Ortholog prediction files were downloaded from the source databases when available. Standalone programs were run locally using the longest protein isoform per gene from the NCBI RefSeq Protein annotation for algorithms for which prediction files were not available (Sup Table 1) (Pruitt and Maglott 2001; Chen *et al*. 2006; Deluca *et al*. 2006; Li *et al*. 2006; Bruford *et al*. 2008; Vilella *et al*. 2009; Lechner *et al*. 2011; Park *et al*. 2011; Mi *et al*. 2016; Kaduk *et al*. 2017; Emms and Kelly 2019; Hu and Friedberg 2019; Nevers *et al*. 2019; Persson *et al*. 2019; Yates *et al*. 2021; Bradford *et al*. 2022; Fuentes *et al*. 2022; Kaldunski *et al*. 2022; Persson and Sonnhammer 2023; Altenhoff *et al*. 2024; Cosentino *et al*. 2024; Oh *et al*. 2025; Tegenfeldt *et al*. 2025; Hernandez-Plaza *et al*. 2026). Even though the longest isoform can reflect technical artifacts in annotations, or comprise rare or aberrant transcripts, using the longest isoform as a representative is still common practice because it is simple, computationally efficient, annotation independent and highly reproducible. DIOPT Arthropods Plus was assembled using only local runs with standalone programs since many of species included are not covered by the resources we integrated.

**Table 1.** Summary of updates to DIOPT in 2011 to 2025.

| DIOPT release | Species covered | Algorithms included | Comments and UI Features |
| --- | --- | --- | --- |
| **1**<br>**(2011)** | 6 (D.mel; H.sap; S.cer; D.rer; M.mus; C.ele) | **9** | Provided ortholog search with DIOPT score; provided pairwise alignment with domain annotation |
| **2**<br>**(2012)** | 6 (D.mel; H.sap; S.cer; D.rer; M.mus; C.ele) | **9** | Same UI as vs1 |
| **3**<br>**(2013)** | 6 (D.mel; H.sap; S.cer; D.rer; M.mus; C.ele) | **10** | Same UI as vs1 |
| **4**<br>**(2014)** | 8 (D.mel; H.sap; S.cer; D.rer; M.mus; C.ele; **S.pom; X.tro**) | **10** | Added two species; same UI as vs1 |
| **5**<br>**(2015)** | 9 (D.mel; H.sap; S.cer; D.rer; M.mus; C.ele; S.pom; X.tro; **R.nor**) | **13** | Added one species; did UI face-lifting; added link-out to UP-TOR for in vivo or in vitro RNAi reagents. |
| **6**<br>**(2016)** | 9 (D.mel; H.sap; S.cer; D.rer; M.mus; C.ele; S.pom; X.tro; R.nor) | **14** | Added paralog search; added 1 to all species ortholog search with heatmap, multi-sequence alignment, link-out to G2F |
| **7**<br>**(2018)** | 10 (D.mel; H.sap; S.cer; D.rer; M.mus; C.ele; S.pom; X.tro; R.nor; **A.tha**) | **17** | Added one species; added user submission and feedback |
| **8**<br>**(2019)** | 10 (D.mel; H.sap; S.cer; D.rer; M.mus; C.ele; S.pom; X.tro; R.nor; A.tha) | **17** | Added API access |
| **9**<br>**(2020)** | 12 (D.mel; H.sap; S.cer; D.rer; M.mus; C.ele; S.pom; X.tro; R.nor; A.tha; **E.coli; A.gam**) | **23** | Added 2 species; did web redesign; added link-out to DRscDB and HPA; supported search of both vs 8 and 9 |
| **10**<br>**(2025)** | 13 (D.mel; H.sap; S.cer; D.rer; M.mus; C.ele; S.pom; X.tro; R.nor; A.tha; E.coli; A.gam; **I.sca**) | **19** | Added one species; removed outdated algorithms; supported search of both vs 9 and 10. |

Red font is used when the species was newly introduced in this release.

An in-house pipeline was developed first to harmonize the outputs from various sources/tools by extracting the protein/gene pairs and synchronizing the protein/gene identifiers to NCBI Gene identifiers. Next, it integrates the harmonized ortholog prediction files from individual resources into one file and identifies the unique gene pairs while counts how many algorithms support each gene pair, generating what we refer to as a DIOPT score. Then, DIOPT pipeline annotates the rank by evaluating for a given gene pair (e.g., A-B) if the score is the highest possible given a forward search (A as input) and/or a reverse search (B as input). A high confidence rank was assigned when the DIOPT score is the highest in both forward and reverse searches and the prediction is supported by at least two algorithms. A moderate confidence rank was assigned when the score is the highest in either the forward or the reverse search but not both, and the pair is supported by at least two algorithms. Moderate confidence was also applied for pairs for which the score is 4 or higher but the pair was not the highest scoring result for either the forward or the reverse search. All other pairs were annotated as low confidence. Many algorithms, such as Compara, OMA and eggNOG, output the prediction in clusters, such that gene pairs outputted by these algorithms include the orthologous relationships across species as well as the paralogous relationships (in-paralogs and outer paralogs).

Pair-wise protein alignments were pre-computed using the Smith-Waterman algorithm using the longest isoform representative per gene based on NCBI RefSeq Protein annotations, and protein domain annotations were obtained from the NCBI Conserved Domain Database (CDD) (Marchler-Bauer *et al*. 2015).

### 2. Ortholist assembly and GSEA analysis

*D. melanogaster*-human orthologous mapping was extracted from the current DIOPT (version 10) and each of the past releases (versions 1 – 9). The NCBI Gene IDs from each release were synchronized to NCBI gene records using a gene history file downloaded from https://ftp.ncbi.nlm.nih.gov/gene/DATA/gene_history.gz on Dec 12^th^ 2025, and the synchronized files were merged. For each fly-human ortholog pair, statistics such as the counts of high, moderate, and low rank pairs across all releases were generated, then an aggregation score was calculated (Count_high_Weight_high_+ Count_moderate_Weight_moderate_+ Count_low_Weight_low_). The weight varies for different ranks: 100 for high rank, 10 for moderate rank and 1 for low rank from version 1-9. The weights were doubled for the current version.

To determine an appropriate aggregation score cutoff, we examined how varying this threshold from 10 to 100 affected several metrics. Specifically, we plotted on the y-axis: (A) the total number of conserved genes, (B) the percentage of conserved genes with fewer than five orthologs, and (C) the average number of orthologs per gene in human and fly, against the aggregation score cutoff on the x-axis (Sup Figure 1). Lower cutoffs yielded more conserved genes (greater gene coverage) but also increased the number of ortholog candidates per gene, as indicated by a lower percentage of genes with fewer than five ortholog candidates and a higher average number of orthologs per gene. This illustrates the trade-off between recall and precision. To balance these two aspects, we selected an aggregation score cutoff of 50 for fly–human orthologous mapping in OrthoList. GSEA analyses were done using PANGEA, which allows the user to input a gene list online and select gene sets, then uses standard over-representation analysis based on the hypergeometric distribution (Hu *et al*. 2023). We used the subset of *Drosophila* genes on Ortholist that have disease-related human orthologs as the input and selected the gene functional group annotation from GLAD (Gene List Annotation for *Drosophila*) (Hu *et al*. 2015) as well as the FlyBase metabolic pathway annotation and phenotype annotation for the GSEA analysis at PANGEA. The background gene set used for GSEA was all *Drosophila* protein-coding genes (13,968).

### 3. Building the standalone pipeline

Programs were developed to process public ortholog prediction results submitted by ortholog prediction groups to Quest for Orthologs (QfO) Orthology Benchmarking Service site (https://orthology.benchmarkservice.org/proxy/projects/2022/) (Nevers *et al*. 2022) based on the EMBL-EBI Reference proteomics annotation (https://www.ebi.ac.uk/reference_proteomes/) (Uniprot 2025). The programs developed include parsers for each prediction result file as well as a program to integrate the results, annotate the results, and map proteins to NCBI Gene Identifiers. The programs were written in Java and are available at GitHub (https://github.com/yanhuihu-code/DIOPT_QFO_pipeline). Users can download and run the DIOPT standalone pipeline locally.

### 4. Implementation of DIOPT Arthropod Plus

The DIOPT Arthropod Plus online tool was built with the Flask framework and it is served via a Gunicorn WSGI instance. The integrative orthologous/paralogous relationships, protein sequence/annotation, and pair-wise alignment results are stored in a MySQL database. The backend was implemented in Python; the frontend uses HTML templates rendered with Twig. JavaScript functionality is provided primarily through jQuery, which is used for AJAX requests and DOM manipulation. Bootstrap is used for layout and styling, and Font Awesome for project iconography. Toast notifications are employed for user pop-up messages. Both the web application and its database are hosted on the O2 high-performance computing (HPC) cluster at Harvard Medical School, operated by the Research Computing group.

### 5. Curation of Human disease genes

We downloaded a list of human genes that have been linked to rare Mendelian disorders, neurodevelopmental diseases (e.g. autism, intellectual disability, epilepsy) as well as cancer from the following databases. Data was accessed on 2/19/2026. All human gene names and symbols used in these databases were converted to the latest official gene symbol using the HGNC database (https://www.genenames.org/).

Rare Diseases: OMIM(https://www.omim.org/) (Amberger *et al*. 2019)

Neurodevelopmental Diseases: SFARI Gene (https://gene.sfari.org/) (Abrahams *et al*. 2013); SysNDD ( https://sysndd.dbmr.unibe.ch/) (Kochinke *et al*. 2016) and Genes4Epilepsy ( https://github.com/bahlolab/Genes4Epilepsy) (Oliver *et al*. 2023).

Cancer: COSMIC Cancer Gene Census (https://cancer.sanger.ac.uk/census#cl_search) (Sondka *et al*. 2018), OncoKB Cancer Gene List (https://www.oncokb.org/cancer-genes) (Chakravarty *et al*. 2017) and TSGene2 ( https://bioinfo.uth.edu/TSGene/download.cgi) (Zhao *et al*. 2016).

### 6. Availability

Both DIOPT and DIOPT Arthropods Plus databases can be freely accessed without restrictions. The URL for the main DIOPT database is at https://www.flyrnai.org/diopt while the URL for DIOPT Arthropod Plus is at https://www.flyrnai.org/apps/diopt_insect/.

## RESULTS

### 1. Major Improvements to the DIOPT online resource

What is known about the function of a gene in one species is often used to make an informed prediction about the function of its orthologs in other species, a concept that is at the foundation of modern biological and biomedical research. High-confidence identification of orthologs is a prerequisite to using ortholog information to formulate hypotheses on gene function. In 2011, we launched the DRSC Integrative Ortholog Prediction Tool (DIOPT), which combines results from multiple ortholog prediction algorithms (Figure 1A,B). This ‘voting system’ approach provides a more sensitive and specific mapping than could be achieved by any given resource. Indeed, we found that the number of resources that predict a given ortholog pair provides a measure of confidence in the results and display the count as part of the DIOPT output results (DIOPT score) (Hu *et al*. 2011). In addition, DIOPT portal provides flexibility through GUI options to filter results at different levels of stringency based on the vote (DIOPT score) or on the annotation confidence (DIOPT rank). DIOPT search results are displayed as a table that provides links to gene-specific information at species-specific and NCBI databases, and provides users with the option to download the results as a tab-delimited file. DIOPT also displays protein and domain alignments, including percent amino acid identity, for inferred ortholog pairs. This helps users identify the most appropriate matches among multiple possible orthologs.

Since the first release launched in 2011, we have made nine additional major releases in the past 14 years (Table 1). Gene and protein information for research species is updated frequently. This leads to corresponding changes to the ortholog prediction results that are generated by application of an ortholog prediction algorithm. At each release we incorporated new gene and protein information, as well as prediction results from the most current version of each prediction tool. In addition, we gradually added new prediction algorithms (Sup Table 1) and expanded species coverage (Sup Table 2). The number of algorithms has increased from 9 to 23 and the number of species included has increased from 6 species to 13. At the most recent release, we removed tools that had not been recently updated, resulting in use of 19 tools/resources in the current version. In recent updates, we also modernized the user interface and implemented new features to enhance user experience. For example, we made it possible for users to query orthologous genes from all species and added a heatmap visualization option that allows users to quickly evaluate predicted conservation of input genes (Figure 1C). Furthermore, we added a paralog search option (release 6) and also made it possible for users to add missing orthologous relationships as well as provide feedback on any orthologous pair (release 7). Over the years we also added link-outs to other resources such as UP-TORR for RNAi reagents (Hu *et al*. 2013), Gene2Fucntion (Hu *et al*. 2017), HPA (Human Protein Atlas) (Uhlen *et al*. 2015) and DRscDB (DRSC scRNAseq Database) (Hu *et al*. 2021) as well as added an Application Programming Interface (API) (release 8) to allow for programmatic access to ortholog predictions, facilitating more efficient workflows for large-scale queries.

**Figure 1.**
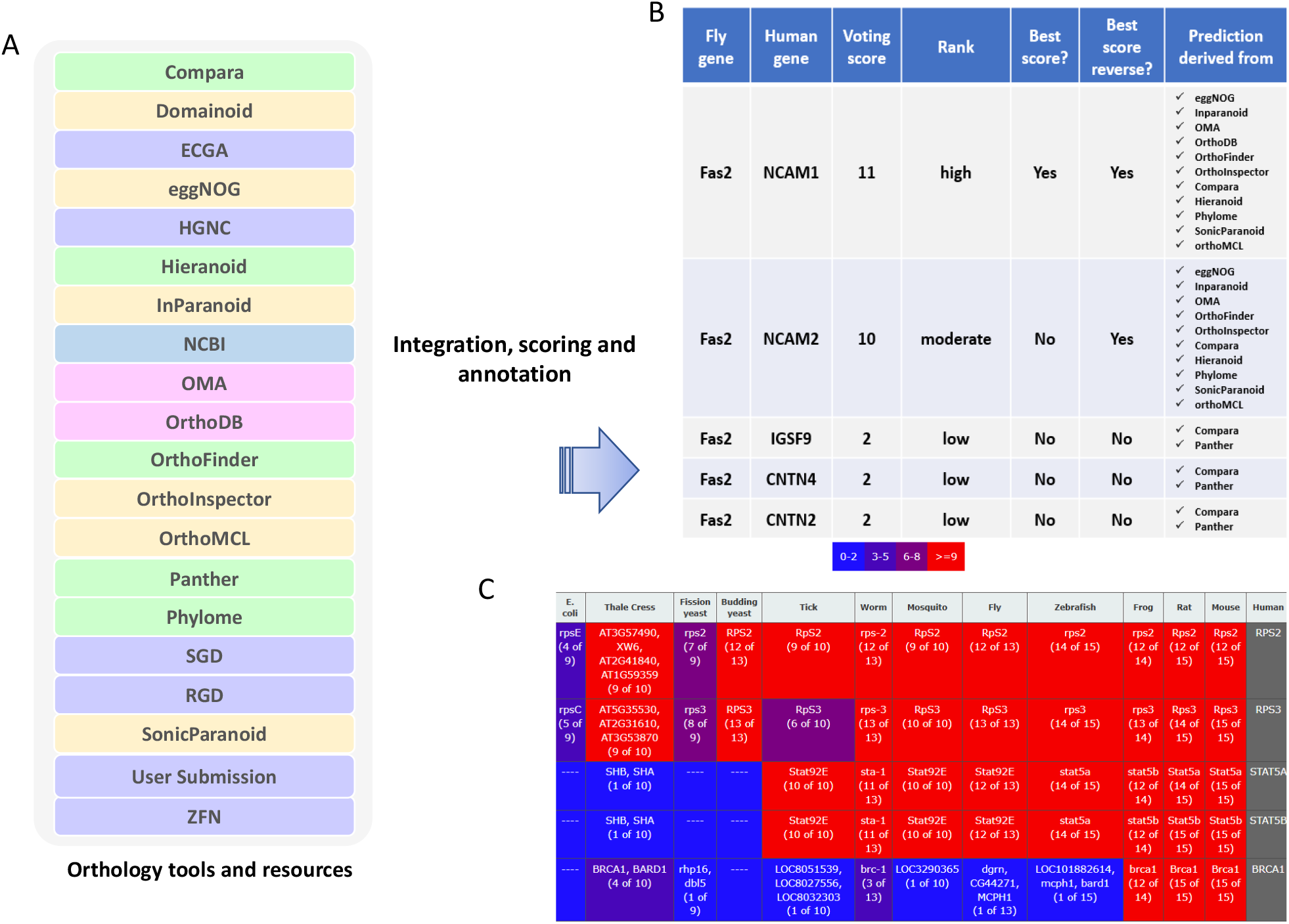
DIOPT, an integrative tool for ortholog and paralog prediction. **A**. DIOPT integrates outputs from several algorithms. Green indicates tree-based algorithms; orange indicates simple graph-based algorithms; the pink indicates evolutionary graph-based algorithms; blue indicates hybrid approaches; purple indicates curated relationships from model organism databases or DIOPT users. **B**. DIOPT results are shown in table format with the voting score (number of tools/resources supporting each pair), an indication if the voting score is the best score in both the forward and reverse searches, and confidence level. **C**. Results of one-to-all searches can be visualized as a heatmap. In this example, human RPS2 and RPS3 are highly conserved across all the species in DIOPT, STAT5A/5B are conserved across animal species, and BRCA1 is mainly conserved in the Tetrapoda species covered in DIOPT.

These changes expanded DIOPT usage to a broad community of researchers interested to identify orthologs as part of studies designed to identify candidate genes to be included in a study, develop new hypotheses regarding variant pathogenicity and/or gene function based on ortholog information, and/or annotate genes based on information from other species.

### 2. Building a Fly-human OrthoList

Genome sequence databases are updated regularly, and each update can lead to revisions in gene and protein annotations. While some of these changes reflect lasting improvements, others may continue to fluctuate as prediction algorithms advance and additional sequencing data become available. For example, annotations of pseudogenes, non-protein-coding genes and protein-coding genes might change back and forth when more datasets become available. Because the quality of ortholog predictions depends on the quality of gene models and protein annotations, ortholog predictions are inherently influenced by the evolving nature of genome annotations.

We compared ortholog mapping between *Drosophila melanogaster* and human genes across multiple DIOPT releases spanning 14 years. Even for well-annotated genomes such as the *Drosophila* and human genomes, approximately 2% of orthologous relationships were dropped in each release, primarily due to withdrawal of gene records or reannotation of gene types from protein-coding to pseudogenes or other non–protein-coding categories. As protein annotations and underlying algorithms have improved over time, only about 25,000 (87%) of the relationships identified in the initial DIOPT release are retained in the current version, while approximately 110,000 new orthologous relationships between *Drosophila* and human genes have been added relative to the first release.

The online DIOPT tool was designed for mapping of orthologs based on user-defined inputs. A genome-wide ortholog mappings, is useful for cross-species comparisons and integration of ‘omics datasets (Hu *et al*. 2018). Two versions of OrthoList, the reference mapping between nematodes and human, have been built in the past by another group (Shaye and Greenwald 2011; Kim *et al*. 2018). To support genome-wide comparisons with *Drosophila*, we constructed a reference OrthoList for fly-human gene mapping by systematically analyzing orthologous predictions from the past 10 DIOPT releases. DIOPT ranks from all releases were incorporated to compute an aggregation score, which was used to select the subset of fly-human ortholog mappings that are consistently present across multiple DIOPT releases. A high confidence rank was annotated when the DIOPT score is the highest at both forward and reverse searches while a moderate confidence rank was annotated when the score is the highest at either forward or reverse search but not both. The high confidence rank is stringent, which works well when a user is seeking one-to-one mapping, although it is useful to include moderate rank pairs when the two species being compared are evolutionary further apart, such as when mapping the simpler

*Drosophila* genome to the human genome. Based on the analysis of gene coverage, the average number of orthologs per gene, and the ratio of genes with a low number of orthologs (Sup Figure 1), the aggregation score cutoff of 50 was chosen to provide a balanced trade-off between recall and precision. Using this cutoff, a total of 23,762 orthologous gene pairs were selected, encompassing approximately 13,000 human genes (63%) and 9,700 *Drosophila* genes (69%) (Sup Table 3). A list of human disease-related genes was assembled based on their association with rare Mendelian diseases (OMIM) (Amberger *et al*. 2019), neurodevelopmental diseases (SFARI Genes, SysNDD, Genes4Epilepsy) (Abrahams *et al*. 2013; Kochinke *et al*. 2016; Oliver *et al*. 2023) and/or cancer (COSMIC, OncoKB, TSGene2) (Zhao *et al*. 2016; Chakravarty *et al*. 2017; Sondka *et al*. 2018); this list includes 6,073 protein-coding genes. A total of 4839 (80%) human disease genes are conserved with *Drosophila* based on our OrthoList, consistent with the idea that *Drosophila* is a valuable model system in which to study human genetic diseases. We next performed gene set enrichment analysis (GSEA) on the subset of *Drosophila* genes on the OrtoList that have disease-related human orthologs using PANGEA (Hu *et al*. 2023). The functional groups that are highly over-represented are metabolic, mitochondria, kinases, transporters, and cytoskeletal genes (Figure 2A), and energy metabolism, glycoconjugate metabolism, cofactor/carriers/vitamin metabolism, and carbohydrate metabolism are among the most enriched metabolic pathways (as annotated at FlyBase; Figure 2B). We found that abnormal phenotypes had been associated with mutations in more than 70% of *Drosophila* orthologs of human disease genes based on FlyBase annotation. Therefore, we also performed GSEA using the gene set based on the FlyBase phenotype annotation (classical alleles); the most enriched phenotypes include physiological phenotypes such as abnormal memory and chemical resistance, and various morphology phenotypes, such as abnormal cell shape or eye color (Figure 2C). To further support collaborative research, we incorporated human disease relevance and additional functional annotations into the FlyOrthoList (Sup Table 3).

**Figure 2.**
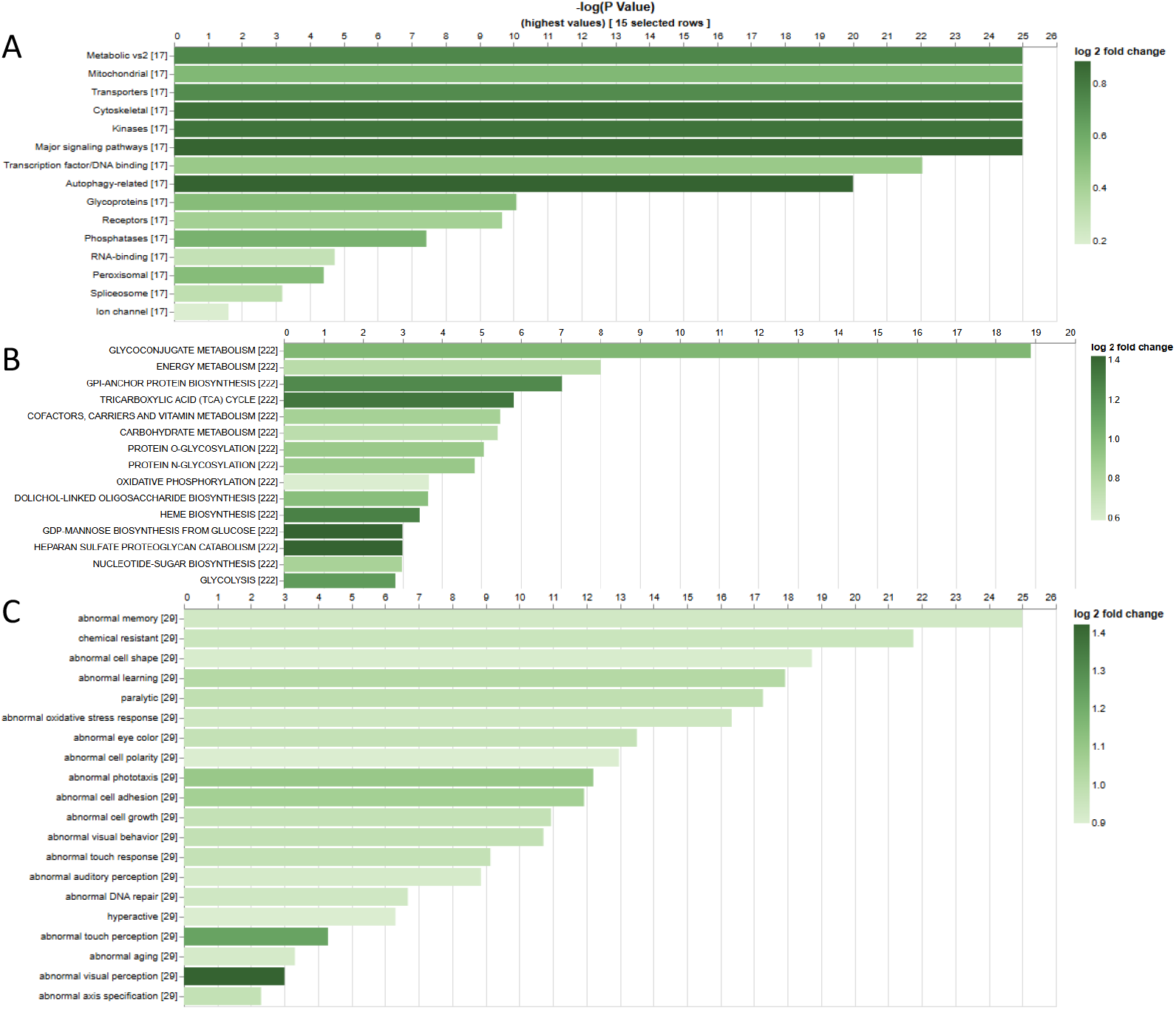
Gene set enrichment analysis of *Drosophila* orthologs of human disease genes. For A, the gene set is the functional group annotations from GLAD (Gene List Annotation for Drosophila). For B, FlyBase metabolic pathway annotations. For C, FlyBase classical alelle phenotype annotations.

### 3. A standalone pipeline for customized assemblies

The first release of DIOPT covers six major model organisms. Over the years, we have received a number of requests from the community to add more species, and we have responded to this by adding seven additional species in subsequent releases. Some of the ortholog prediction algorithms (eg. OMA and orthoDB) output result in cluster format while others (e.g., InParanoid) output predictions in pair format. To integrate these two types of predictions, orthologous or paralogous gene pairs need to be extracted first from each cluster for all the cluster-based algorithms, then pair-wise relationships can be integrated. Although growth is linear when adding more species to a cluster-based tool, however, growth is quadratic for pair-based results. Therefore, scaling up to include more species in a pair-based web tool is challenging because of the significant increase of records in database and the compromise of query performance associated with that increase. For example, there are about 0.771 million gene pairs in DIOPT version 1 for the six species covered and this expanded to 33 million gene pairs in DIOPT version 10 for coverage of 13 species. When the number of species doubled, the number of records in the database table that tracks gene pairs increased more than 41 times, and the number of records in the table tracking the original tools for each gene pair increased from about 2 million to 63 million records. Therefore, it is not practical to add all the species we have been requested to add. Another factor that guided our decision regarding what species to include is that the DIOPT online database is primarily focused on well-studied model organisms. To accommodate requests outside of a core set of species, we implemented a standalone pipeline that integrates predictions from multiple algorithms; specifically, the subset of algorithms at DIOPT that have been submitted by their developers to the Quest for Orthologs (QfO) Orthology Benchmarking Service (Nevers *et al*. 2022). This standalone pipeline can be easily customized by the user to accommodate the species of interest.

Using this pipeline along with the main database, DIOPT-based ortholog mapping outputs have been integrated into several public resources, including FlyBase (Ozturk-Colak *et al*. 2024), the Alliance of Genome Resources (Alliance of Genome Resources 2024), and MARRVEL (Wang *et al*. 2017), to facilitate functional genomic studies. For example, DIOPT has been integrated in MARRVEL (Wang *et al*. 2017), where the orthologous relationships and their protein alignments from DIOPT have been used to evaluate the SNPs from human diseases and identify the appropriate animal models for follow-up studies (Sup Figure 2).

QFO relies on the EMBL-EBI reference proteome which currently covers 81 species (UniProt Release 2025_04), including key model organisms, and prioritizes high-quality, well-annotated genomes that represent major evolutionary lineages (Uniprot 2019). The standalone pipeline relies on precomputed predictions from QFO and therefore is limited to species covered by reference EMBL-EBI reference proteome.

### 4. Implementation of DIOPT Arthropod Plus, a DIOPT instance with a specific focus

The majority of the biological research papers are focused on model organisms that are well annotated based on high-quality sequence assembles and supporting datasets. We continue to keep the main focus of the DIOPT database on well-studied organisms that serve as anchor species for different evolutionary branches. Given our expertise and interest in arthropod biology, we were interested to develop a sister version of DIOPT that facilitates use of *Drosophila* and other information to inform understanding of other arthropods. Broader motivation for this effort includes the relevance of some arthropods to human and animal health (e.g., mosquito and tick vectors of diseases), as crop pests (e.g., cotton bollworms, corn earworms, and fall armyworms), or as beneficial species (e.g., honeybees and silkworms). Despite the significance of these species, there is a lack of comprehensive informatics tools to aid research. To address this, we launched DIOPT Arthropod Plus, which allows users to query ortholog mapping across multiple non-model arthropod species and key model organisms, including *Drosophila, C. elegans*, and human (Figure3, Sup Table2). At DIOPT Arthropod Plus, users can choose an input and output species from the dropdown menu, then enter a list of genes. Filters at different stringency levels based on voting scores or confidence ranks are provided. A summary of ortholog relationships when output species is different from input species, or of paralogous relationships when input and output species are the same, is provided at the results page. The results page output includes the relevant genes, voting scores, and annotations indicating if the voting score is the best score in the forward and/or reverse direction, as well as a confidence level determined based on these metrics. Users have the option to choose which columns are displayed and download the result as a table. For each inferred ortholog pair, a link is provided to a protein alignment page with a pair-wise protein alignment generated using the Smith-Waterman algorithm. Only the longest protein isoform is selected to represent each gene, and protein domain annotation are highlighted on the aligned sequences. In addition, alignment statistics, such as percent identity, are displayed for both the overall aligned sequences and the protein domain regions (Figure 3).

**Figure 3.**
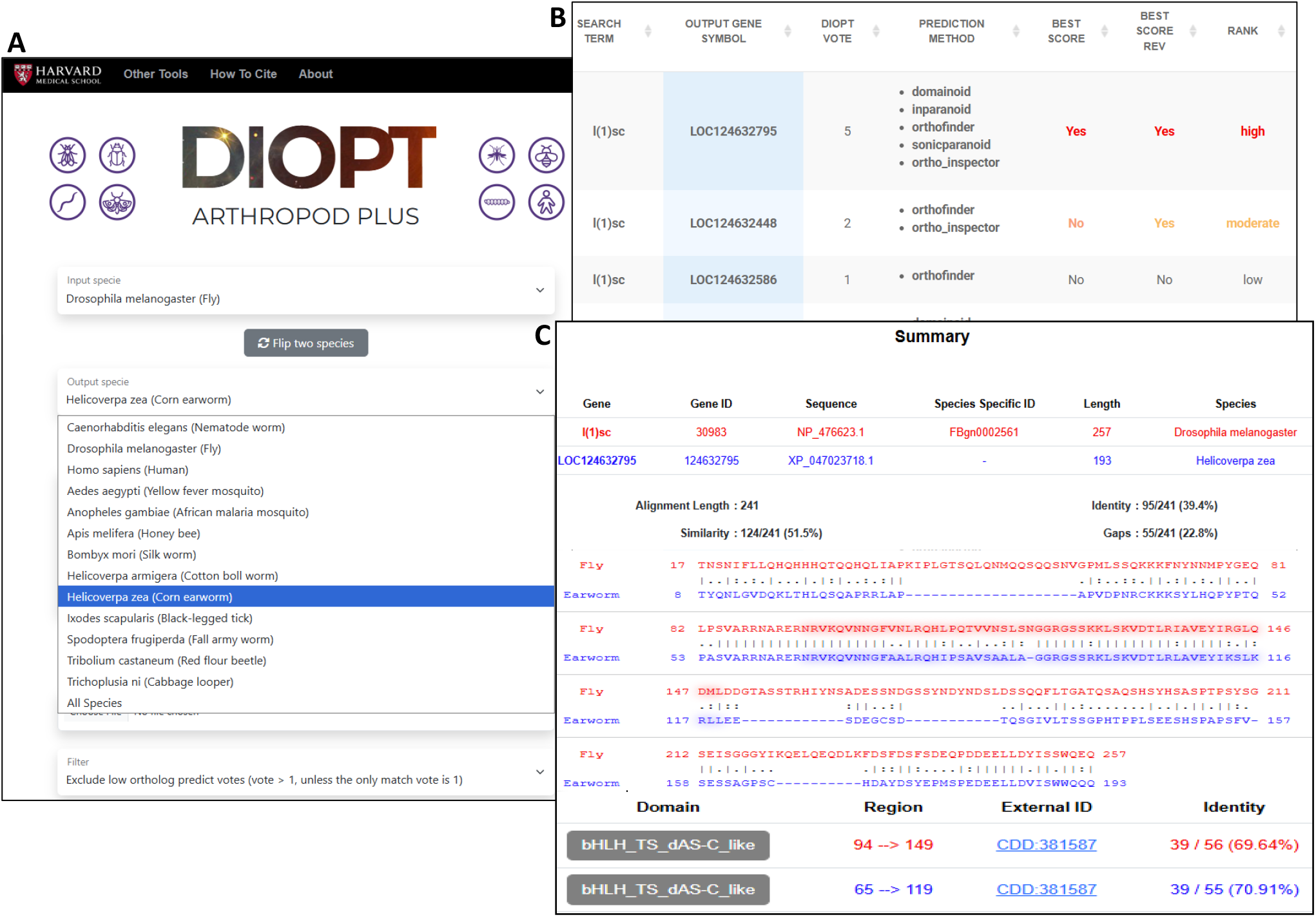
UI features of DIOPT Arthropod Plus. **A**. The search page allows users to choose an input and output species from a menu of 13 species, then enter a list of genes from the input species. Filters facilitate searches at different levels of stringency. **B**. Ortholog or paralog output tables provide a voting score, a list of algorithms that predict the pair, an indication if the score is the best score in both the forward and reverse directions, and an overall confidence score. **C**. Pairwise protein alignment is provided for the longest protein isoforms, and includes summary statistics for amino acid similary and identity, and gaps in the alignment. Functional domains are highlighted on the alignment and the identities over protein domain regions are also indicated.

The DIOPT Arthropod Plus database covers 13 species including 10 insect species (class Insecta) as well as human, *Ixodes scapularis* (black-legged tick), and *C. elegans* genomes (Sup Figure 3). Based on high- and moderate-confidence DIOPT ranks, Noctuidae species (cotton bollworm, corn earworm, fall armyworm, and cabbage looper) show the highest proportion of conserved genes, ranging from 86% to 94%. Approximately 46–49% of human genes are conserved in the various insect species. *Drosophila* shares about 60–67% of its genes with other insects. In contrast, only 29–31% of *C. elegans* genes are conserved with insect species (Figure 4). Our analysis confirmed that a larger fraction of fly genes (67%) has clear orthologs in other insects (including Noctuidae, mosquitoes, beetles, etc.) due to the closer evolutionary distance. Consequently, *Drosophila* can serve as a “reference insect” for interpreting data from non-model species (Mameli *et al*. 2025) and for improving gene prediction and functional annotations in their genomes. DIOPT Arthropod Plus will enable researchers to rapidly generate informed hypotheses regarding gene functions and interactions that would be difficult to characterize first in non-model species.

**Figure 4.**
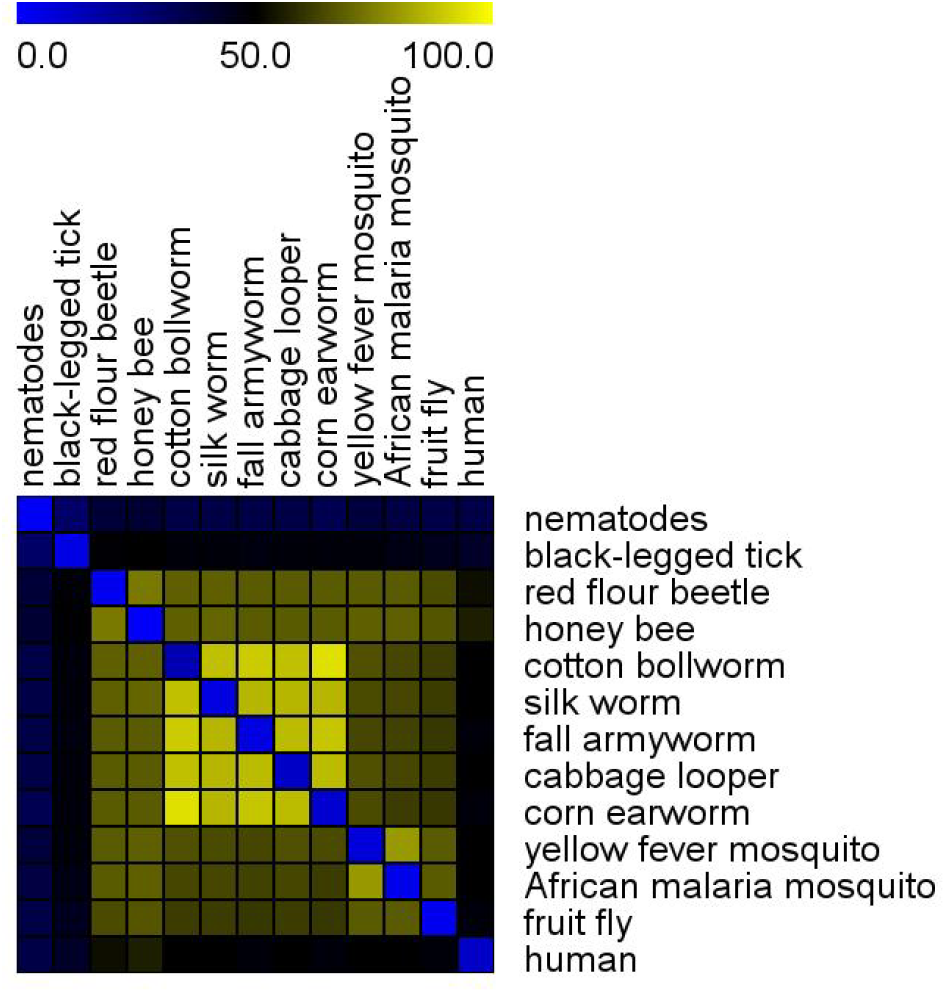
Conservation among species included in DIOPT Arthropod Plus. Shown is a heatmap of the percentage of conserved genes between any two species included in the resource, using a rank of high or moderate as the cutoff.

## DISCUSSION/SUMMARY

We describe the decades-long effort to maintain and improve DIOPT, an integrative ortholog mapping resource. Major changes made in the ten DIOPT versions that have been released since the initial publication in 2011 are summarized in Table 1. In addition, *Drosophila-*human ortholog relationships from versions 1 to 10 have been analyzed to build FlyOrthoList (Sup Table 3), a reference ortholog mapping between fly and human that covers about 9,700 of *Drosophila* genes (69% of *Drosophila* protein-coding genes) and 13,000 human genes (63% of human protein-coding genes). Of the 6,073 human protein-coding genes associated with diseases, 4,839 (80%) are conserved in *Drosophila*; moreover, 77% of these fly genes are associated with mutant phenotype annotations. The rich research resources available for *Drosophila*, such as mutant collections, transgenics, CRISPR tools, cloned cell-type specific drivers/enhancers, and extensive functional annotations, make *Drosophila* a powerful model organism for studying human diseases.

In addition, we further expanded DIOPT by creating a pipeline useful for DIOPT-based ortholog mapping for user-inputted species, as well as a sister database, DIOPT Arthropod Plus, which was implemented to facilitate research in non-model arthropods relevant to human health and food security. Altogether, the DIOPT suite of resources comprise a set of tools that can be used without specialized understanding of bioinformatics to support functional genomics studies.

## ACKNOWLEDGEMENTS

We thank the past and current members from Perrimon lab, DRSC and TRiP for their valuable suggestions and feedbacks. We extend our gratitude to the Harvard Medical School Research Computing and IT-Client Services teams for their consultation, web hosting, and support.

The article submitted together with this notice is subject to the Immediate Access to Research policy of the Howard Hughes Medical Institute (“HHMI”). In accordance with this policy: (i) a preprint of this article either has been, or will be, deposited on a preprint server under a Creative Commons Attribution 4.0 International (CC BY 4.0) license and (ii) an additional author-published revised version of this article incorporating peer review feedback and/or new results or analysis either has been, or prior to journal publication will be, deposited on a preprint server under a CC BY 4.0 license. In addition, a non-exclusive CC BY 4.0 license to this article has been granted to the public and HHMI has a sublicensable, non-exclusive license to this article.

## DATA AVAILABILITY STATEMENT

The resources are available to public. The URL for DIOPT is https://www.flyrnai.org/diopt and the URL for DIOPT Arthropods plus is www.flyrnai.org/apps/diopt_insect/. The standalone pipeline is available at https://github.com/yanhuihu-code/DIOPT_QFO_pipeline.

