## Supplemental Figures for "DIOPT: the DRSC Integrative Ortholog Prediction Tool, 2026 update"

**Sup Figure 1. Metrics used to set the cutoff of robustness score for FlyOrthoList**

To determine an appropriate aggregation score cutoff, we examined how varying this threshold from 10 to 100 affected several metrics. Specifically, we plotted on the y-axis: (A) the total number of conserved genes, (B) the percentage of conserved genes with fewer than five orthologs, and (C) the average number of orthologous genes per gene in human and fly, respectively, against the aggregation score cutoff on the x-axis.

Lower cutoffs yielded more conserved genes (greater gene coverage) but also increased the number of ortholog candidates per gene, as indicated by a lower percentage of genes with fewer than five ortholog candidates and a higher average number of orthologs per gene. This illustrates the trade-off between recall and precision. To balance these two aspects, we selected an aggregation score cutoff of 50 for fly–human orthologous mapping in OrthoList.

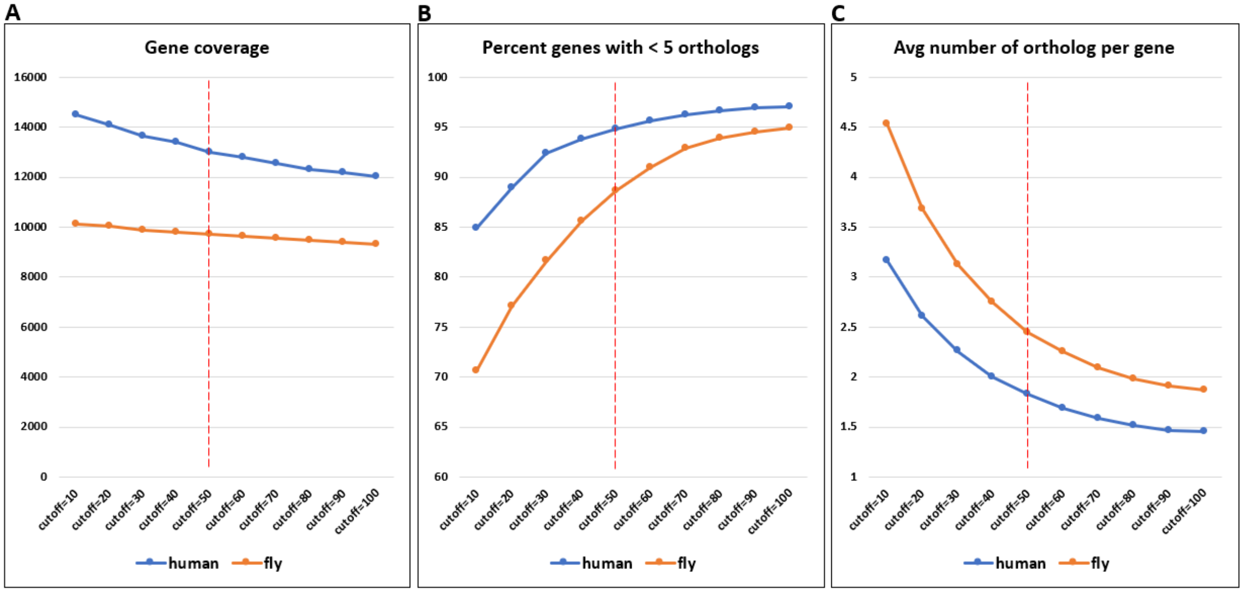

**Sup Figure 2. DIOPT supports ortholog mapping at many public resources.** Either DIOPT database exports or results from the customizable DIOPT pipeline have been integrated into the indicated public resources.

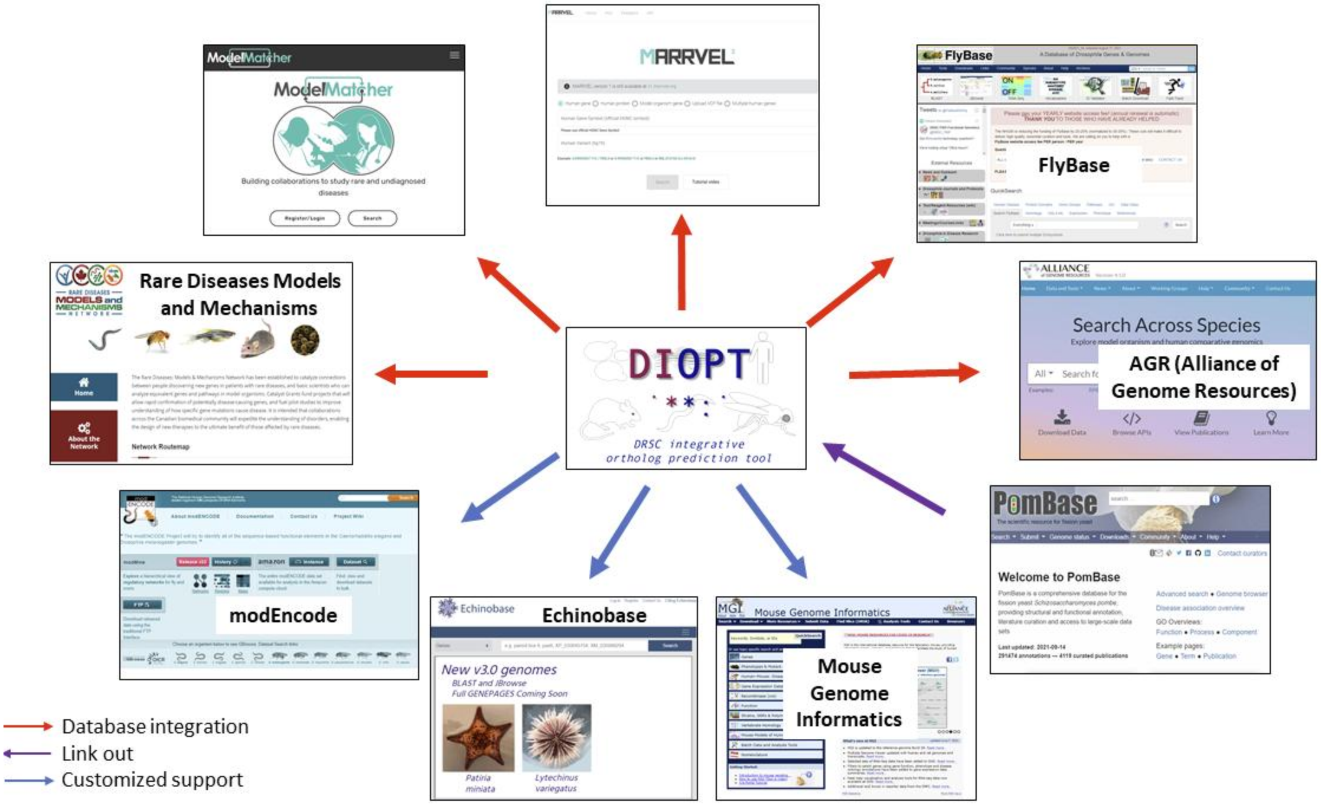

### Sup Figure 3. Species in DIOPT arthropod plus

```
Bilateria
├── Deuterostomia
│   └── Chordata
│       └── Mammalia
│           └── Primates
│               └── Homo sapiens
└── Protostomia
    └── Ecdysozoa
        ├── Nematoda
        │   └── Caenorhabditis elegans
        └── Arthropoda
            ├── Chelicerata
            │   └── Ixodida (ticks)
            │       └── Ixodes scapularis
            └── Hexapoda (Insecta)
                ├── Hymenoptera
                │   └── Apis mellifera
                ├── Coleoptera
                │   └── Tribolium castaneum
                ├── Diptera
                │   ├── Drosophilidae
                │   │   └── Drosophila melanogaster
                │   └── Culicidae (mosquitoes)
                │       ├── Aedes aegypti
                │       └── Anopheles gambiae
                └── Lepidoptera
                    ├── Bombycidae
                    │   └── Bombyx mori
                    └── Noctuidae
                        ├── Helicoverpa armigera
                        ├── Helicoverpa zea
                        ├── Spodoptera frugiperda
                        └── Trichoplusia ni
```
